# Characterization of the composition and functioning of the Crumbs complex in *C. elegans*

**DOI:** 10.1101/2021.08.10.455623

**Authors:** Victoria G Castiglioni, João J Ramalho, Jason Kroll, Riccardo Stucchi, Hanna van Beuzekom, Ruben Schmidt, Maarten Altelaar, Mike Boxem

## Abstract

The apical domain of epithelial cells can acquire a diverse array of morphologies and functions, which is critical for the function of epithelial tissues. The Crumbs proteins are evolutionary conserved transmembrane proteins with essential roles in promoting apical domain formation in epithelial cells. The short intracellular tail of Crumbs proteins interacts with a variety of proteins, including the scaffolding protein Pals1 (protein associated with LIN7, Stardust in *Drosophila*). Pals1 in turn binds to a second scaffolding protein termed PATJ (Pals1-associated tight junction protein), to form the core Crumbs/Pals1/PATJ Crumbs complex. While essential roles in epithelial organization have been shown for Crumbs proteins in *Drosophila* and mammalian systems, the three *Caenorhabditis elegans crumbs* genes are dispensable for epithelial polarization and animal development. Moreover, the presence and functioning of orthologs of Pals1 and PATJ has not been investigated. Here, we identify MAGU-2 and MPZ-1 as the *C. elegans* orthologs of Pals1 and PATJ, respectively. We show that MAGU-2 interacts with all three Crumbs proteins as well as MPZ-1, and localizes to the apical membrane domain in a Crumbs-dependent fashion. Similar to *crumbs* mutants, a *magu-2* null mutant shows no developmental or epithelial polarity defects. Finally, we show that overexpression of the Crumbs proteins EAT-20 or CRB-3 in the *C. elegans* intestine can lead to apical membrane expansion. Our results shed light into the composition of the *C. elegans* Crumbs complex and indicate that the role of Crumbs proteins in promoting apical domain formation is conserved.

## Introduction

Cell polarity, the asymmetric distribution of components and functions in a cell, is a fundamental property of animal cells. In epithelial cells, proteins and lipids of the plasma membrane are distributed asymmetrically into an apical domain that faces the external environment or lumen and a basolateral domain contacting neighbouring cells and the extracellular matrix. Cell-cell junctions at the boundary of the apical and basolateral domains seal the epithelial sheets to segregate the internal medium from the outside environment and provide mechanical strength to the tissue. The establishment of these different domains relies on mutually antagonistic interactions between evolutionarily conserved polarity complexes. Members of the Par and Crumbs complexes establish apical identity and position cell junctions at the apical/lateral border, while basolateral identity is promoted by the Scribble group proteins and the Par1 kinase (Rodriguez-Boulan and Macara, 2014; Wen and Zhang, 2018).

The Crumbs complex is a key regulator of epithelial polarity that is crucial for the formation of the apical domain and the organization of junctions, and thus the boundary between the apical and basolateral domains (Klebes and Knust, 2000; Médina et al., 2002b; Tepass et al., 1990; Wodarz et al., 1995). There are multiple ways in which the Crumbs complex promotes epithelial polarity. For example, direct interactions between Crumbs and proteins that bind the actin cytoskeleton such as Moesin support the polarized organization of the cytocortex (Médina et al., 2002a). Crumbs can also recruit β_Heavy_-Spectrin to the apical membrane, contributing to apical membrane stability (Lee and Thomas, 2011; Pellikka et al., 2002), and promotes the apical localization and accumulation of the apical Par proteins (Almeida et al., 2019; Lemmers et al., 2004). Loss of the Crumbs complex leads to severe polarity defects (Straight et al., 2004; Wodarz et al., 1993), disruption of cell junctions (Shin et al., 2005; Wang et al., 2006) and loss of tissue integrity (Jürgens et al., 1984). Nevertheless, the Crumbs complex is not essential in all epithelial tissues. For example, the *Drosophila* intestine and neuroblasts can withstand the absence of the Crumbs complex without apparent defects (Hong et al., 2001; Tepass et al., 1990).

The Crumbs complex is composed of three core scaffolding proteins: the transmembrane protein Crumbs (Crb in *Drosophila;* CRB1, CRB2, and CRB3 in mammals), the MAGUK (membrane-associated guanylate kinase) protein Pals1 (Protein associated with LIN7, Sdt in *Drosophila)*, and Pals1-associated tight junction protein (PATJ) (Bhat et al., 1999; Knust et al., 1993; Nam and Choi, 2006; Tepass and Knust, 1993). Whereas *Drosophila* encodes one Crumbs protein, several orthologs can be found in the *C. elegans*, zebrafish, mouse, and human genomes. Mammalian CRB1 and CRB2 are similar in domain composition to *Drosophila* Crumbs. They are composed of a large extracellular region, a transmembrane domain, and a small highly conserved intracellular region containing two protein-protein interaction domains: a band 4.1/Ezrin/ Radixin/Moesin (FERM)-binding motif (FBM) and a C-terminal “ERLI” PDZ-binding motif (PBM). CRB3 is similar to CRB1 and CRB2 but lacks the large extracellular region. Pals1 contains two Lin-2 and Lin-7 (L27) domains, a postsynaptic density 95/disks large/zona occludens-1 (PDZ) domain, a Src homology 3 (SH3) domain and a guanylate kinase (GUK) domain, whereas PATJ contains one N-terminal L27 domain and multiple PDZ domains (Assémat et al., 2008).

To form the Crumbs complex, Pals1 interacts with the PBM of Crumbs through its PDZ and SH3 domains, additionally strengthening the interaction through its GUK domain (Bachmann et al., 2001; Hong et al., 2001). Pals1 and Patj form an L27 tetramer through the interaction between two units of Pals1-L27N— PATJ-L27 (Feng et al., 2005; Roh et al., 2002). The core Crumbs complex can recruit additional proteins, like Moesin, Lin-7, or Par-6, depending on the requirements of the cell (Bulgakova and Knust, 2009). Moreover, Crumbs and PATJ can form an alternative complex through interactions with Par-6 rather than Pals1.

Like mammals, *C. elegans* encodes three Crumbs proteins, CRB-1, EAT-20, and CRB-3, that are similar in domain composition to their mammalian counterparts (Bossinger et al., 2001; Waaijers et al., 2015). The subcellular localization of the Crumbs proteins is strikingly similar to that of Crumbs proteins in mammalian systems and *Drosophila* (Bossinger et al., 2001; Waaijers et al., 2015). CRB-1 can provide a positional cue for junction formation in the intestine of animals depleted of both HMP-1 α-catenin and LET-413 Scribble (Segbert et al., 2004). However, *C. elegans* Crumbs proteins are not essential for epithelial development in *C. elegans*, and even triple *crumbs* deletion mutants are viable without obvious defects in cell polarity (Bossinger et al., 2001; Segbert et al., 2004; Waaijers et al., 2015). Thus, the precise role of the Crumbs complex in *C. elegans* remains elusive. In addition, the composition of the Crumbs complex in *C. elegans* has not been investigated. Even though putative orthologs of Pals1 and PATJ exist, it is not known whether they interact with each other or with any of the Crumbs proteins.

Here, we investigate whether the composition of the Crumbs complex is conserved in *C. elegans*. Additionally, we investigate if the Crumbs proteins can promote apical domain formation through overexpression studies. We demonstrate that MAGU-2 interacts with the three Crumbs proteins and is localized to the apical membrane domain in a Crumbs-dependent manner. Moreover, we find that MAGU-2 can interact with the putative PATJ ortholog MPZ-1. Thus, our data strongly suggests that MAGU-2 is the *C. elegans* homolog of Pals1 and a member of the Crumbs complex. Like triple *crumbs* deletion mutants, animals lacking *magu-2* are viable and show no overt polarity defects. However, overexpression of EAT-20 and CRB-3 in *C. elegans* resulted in apical membrane expansion, as reported for overexpression of *Drosophila* Crumbs. Our results therefore indicate that the composition of the Crumbs complex and the role of Crumbs proteins in apical domain formation are conserved in *C. elegans*.

## Results

### MAGU-2 is the *C. elegans* ortholog of Pals1 and interacts with Crumbs proteins

To identify candidate Pals1 homologs, we searched the predicted *C. elegans* proteome using BLAST with the human Pals1 and *Drosophila* Sdt sequences. We identified two proteins, MAGU-1 and MAGU-2, that have high sequence similarity, and both contain a PDZ domain, an SH3 domain, and a GUK domain. However, neither protein appears to harbor the L27 domains present in Pals1. Of the two, MAGU-2 was more closely related to Pals1 and Sdt than MAGU-1 (Fig. 1A). Reciprocal BLAST using MAGU-2 furthermore identified human Pals1 and *Drosophila* Sdt as the closest homologs, while BLAST using MAGU-1 identified human MPP7 and *Drosophila* Metro as the highest hits. MAGU-2 has two predicted isoforms, MAGU-2a of 830 amino acids (aa) and MAGU-2b of 668 aa. Both isoforms consist of a PDZ domain, an SH3 domain and a GUK domain, while MAGU-2A contains an extra N-terminal PDZ domain which is not present in Pals1 (Fig. 1B). The PDZ and SH3 domains of human Pals1 are involved in the interaction with Crumbs, while the GUK domain further enhances the binding. *Drosophila* Crumbs, in turn, interacts with Pals1 through its intracellular PDZ-binding domain (Li et al., 2014). Most of the amino acids involved in the interaction between Pals1 and Crumbs are conserved in *C. elegans* MAGU-2 and CRB-1/EAT-20/CRB-3 (Fig. 1C). Given the best reciprocal blast, the conservation at the amino acid level and of the domain structure, we conclude that MAGU-2 is the *C. elegans* ortholog of Pals1.

**Figure 1.**
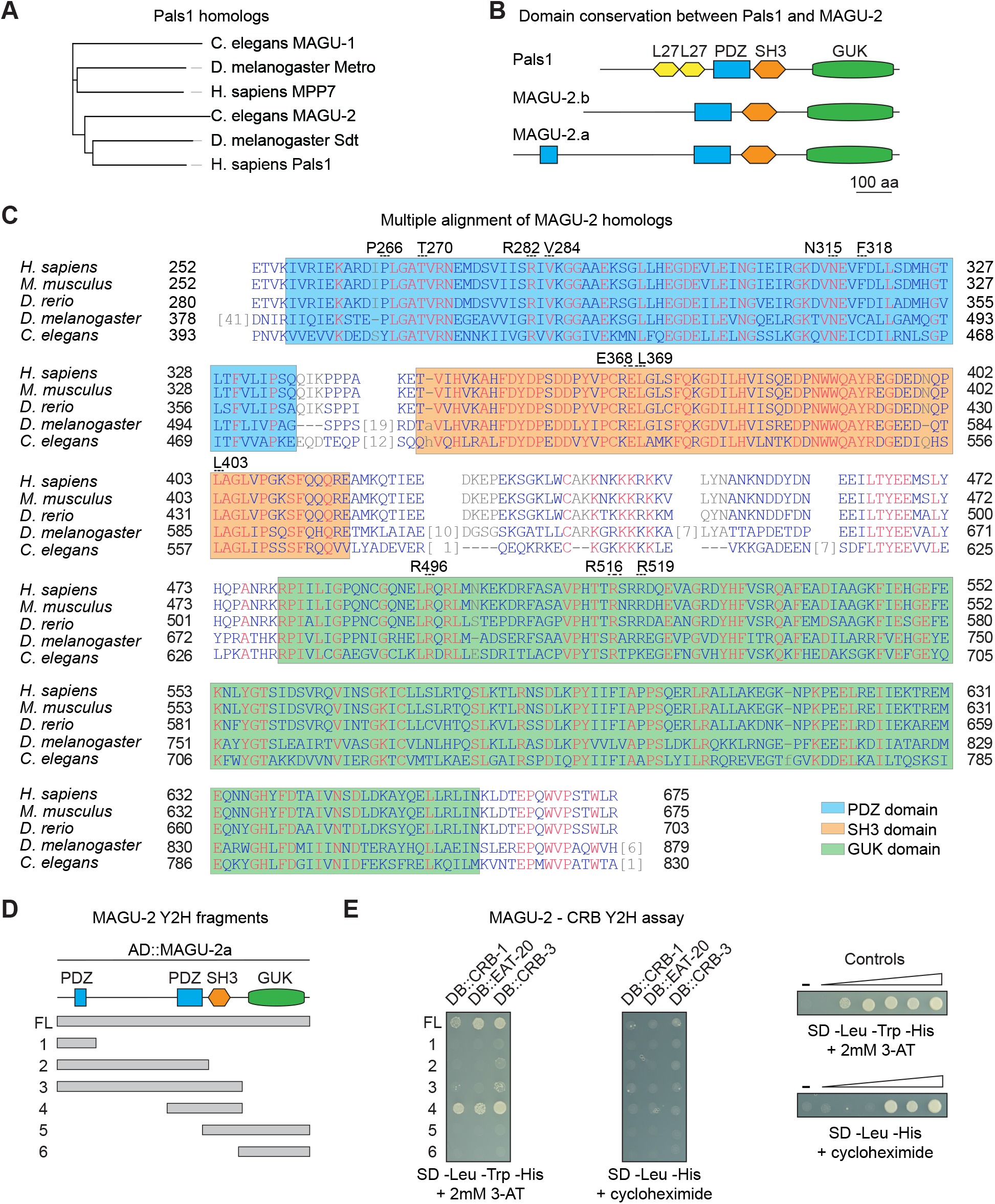
MAGU-2 is the homolog of Pals1/Sdt. **(A)** Phylogenetic tree of Pals1 homologs, including *C. elegans* MAGU-1 and MAGU-2. **(B)** Schematic representation of predicted protein domains of Pals1 and MAGU-2. **(C)** Multiple alignment of MAGU-2a with *Homo Sapiens* Pals1, *Mus musculus* Pals1, *Danio rerio* mpp5a, and *Drosophila melanogaster* Sdt. Numbered residues above the sequence are the Pals1 residues involved in the interaction with Crumbs. **(D)** Schematic representation of the different fragments of MAGU-2 used for the Y2H assay. **(E)** Interaction of MAGU-2 with CRB-1, EAT-20, and CRB-3 in the yeast two-hybrid assay. Fragments of MAGU-2.b fused to the Gal4 activation domain were co-expressed with the intracellular domain of either CRB-1, EAT-20 or CRB-3 fused to the Gal4 DNA binding domain. Growth on the -Leu -Trp -His + 2 mM 3-AT plate indicates presence of interaction. Lack of growth on -Leu -His + cycloheximide plate shows that DB::CRB-1/ EAT-20/CRB-3 are not self-activating. Controls range from no reporter activation to strong reporter activation. Abbreviations: L27, Lin-2 and Lin-7 domain; PDZ, Postsynaptic density 95, Discs large, Zonula occludens-1; SH3 domain, SRC homology 3 domain; GUK, guanylate kinase-like domain.

To determine if MAGU-2 interacts with any of the three *C. elegans* Crumbs proteins, we performed a yeast two hybrid (Y2H) assay. We fused the intracellular domain of the three Crumbs proteins to the DNA binding domain of Gal4, and fused the following fragments of MAGU-2 to the activation domain: a full-length fragment (FL), an N-terminal fragment with the PDZ present in the long MAGU-2 isoform (1), an N-terminal fragment including both PDZ domains but lacking the SH3 domain (2), an N-terminal fragment that includes both PDZ domains and the SH3 domain (3), a fragment containing the second PDZ domain and the SH3 domain (4), a C-terminal fragment containing the SH3 and GUK domains (5), and a C-terminal fragment consisting of only the GUK domain (6) (Fig. 1D). The Y2H assay indicated that full length MAGU-2 interacts with each of the three Crumbs proteins (Fig. 1E). Consistent with previous analyses in other organisms (Li et al., 2014), both the conserved PDZ domain and the SH3 domain are required for the interaction. These results indicate that the interactions between Crumbs and Pals1 are conserved in *C. elegans*, and that MAGU-2 is a member of the *C. elegans* Crumbs complex.

### MAGU-2 localizes apically in epithelial tissues

To determine a potential role for MAGU-2 in organizing epithelial polarity, we first examined its expression pattern and subcellular localization. Mammalian Pals1 and *Drosophila* Std localize at the apical cortex in epithelial tissues. Thus, if it is a functional ortholog, MAGU-2 would be expected to recapitulate the apical localization of these proteins. To visualize the localization of MAGU-2, we used CRISPR/Cas9 to insert the sequence encoding the green fluorescent protein (GFP) into the endogenous *magu-2* locus. The tag was inserted at the C-terminus such that both isoforms are tagged (Fig. 2A). We first detected MAGU-2::GFP in the intestine during early embryonic development, before apical-basal polarity is fully established, together with DLG-1 at the nascent apical domain (Fig. 2B-B’). During later pharyngeal and intestinal development, MAGU-2 localized to the apical cortex, where it colocalized with the apical protein PAR-6 (Fig. 2C–D, C’-D’, F, F’). Throughout larval development, MAGU-2::GFP localized to the nerve ring and to the apical membrane domain of the pharynx and intestinal cells (Fig. 2E–F, E’-F’). The apical subcellular localization of MAGU-2 in polarized epithelial tissues, which matches the subcellular localization of the Crumbs proteins in these tissues (Shibata et al., 2000; Waaijers et al., 2015), strengthens our hypothesis that MAGU-2 is a functional Pals1 ortholog and a member of the Crumbs complex.

**Figure 2.**
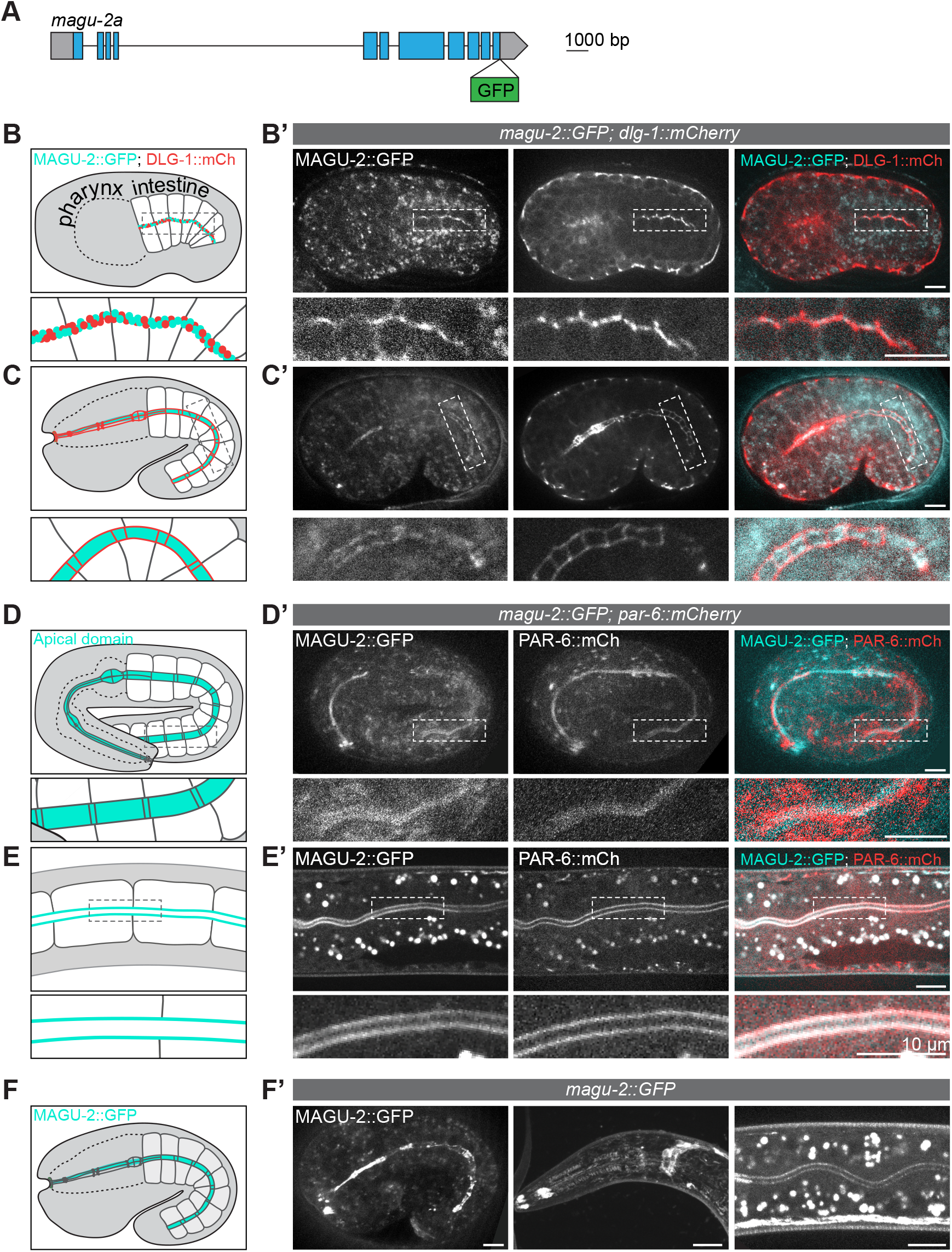
MAGU-2 localizes apically in epithelial tissues. **(A)** Schematic representation of endogenous tagging of *magu-2a* with the sequence encoding a green fluorescent protein (GFP). **(B-F)** Schematic representations of the areas imaged in B’-F’, with the localization of MAGU-2 and PAR-6 indicated in cyan and the localization of DLG-1 indicated in red. **(B’, C’)** Distribution of MAGU-2::GFP and DLG-1::mCherry in the early stages of intestinal polarization and in comma-shape embryos. **(D’, E’)** Distribution of MAGU-2::GFP and PAR-6::mCherry in two-fold embryos and L3 larvae. **(F’)** Distribution of MAGU-2::GFP in the pharynx and intestine of a comma-stage embryo, and the nerve ring and intestine of L2/L3 larvae.

### MAGU-2 is not essential for epithelial polarity

Loss of Pals1 in mammalian and *Drosophila* epithelia leads to severe polarity defects and disruption of cell junctions (Hong et al., 2001; Straight et al., 2004; Wang et al., 2006). The phenotypes of Crumbs and Pals1 mutants are very similar, consistent with them forming a core complex (Hong et al., 2003; Nam and Choi, 2006; Tepass and Knust, 1993). To investigate the role of *magu-2* in polarity establishment we used CRISPR-Cas9 to generate a *magu-2* null allele, *mib6*, that removes 806 out of the 830 amino acids of the long isoform and introduces an early frame shift in the MAGU-2 coding sequence of both isoforms (Fig. 3A). Animals homozygous for *magu-2(mib6)* are viable and do not display any obvious developmental defects, suggesting that MAGU-2 activity is dispensable for *C. elegans* development (Fig. 3B). In order to address any redundancy between MAGU-2 and the Crumbs proteins, we examined the effect of combining the *magu-2* deletion with the triple *crumbs* deletion. We previously reported that a triple *crumbs* deletion results in a small but significant reduction in progeny numbers (Waaijers et al., 2015). Combining the *magu-2* deletion with the triple *crumbs* deletion did not exacerbate the phenotype of the *crumbs* deletion mutant (Fig. 3B), consistent with both proteins acting in the same pathway.

**Figure 3.**
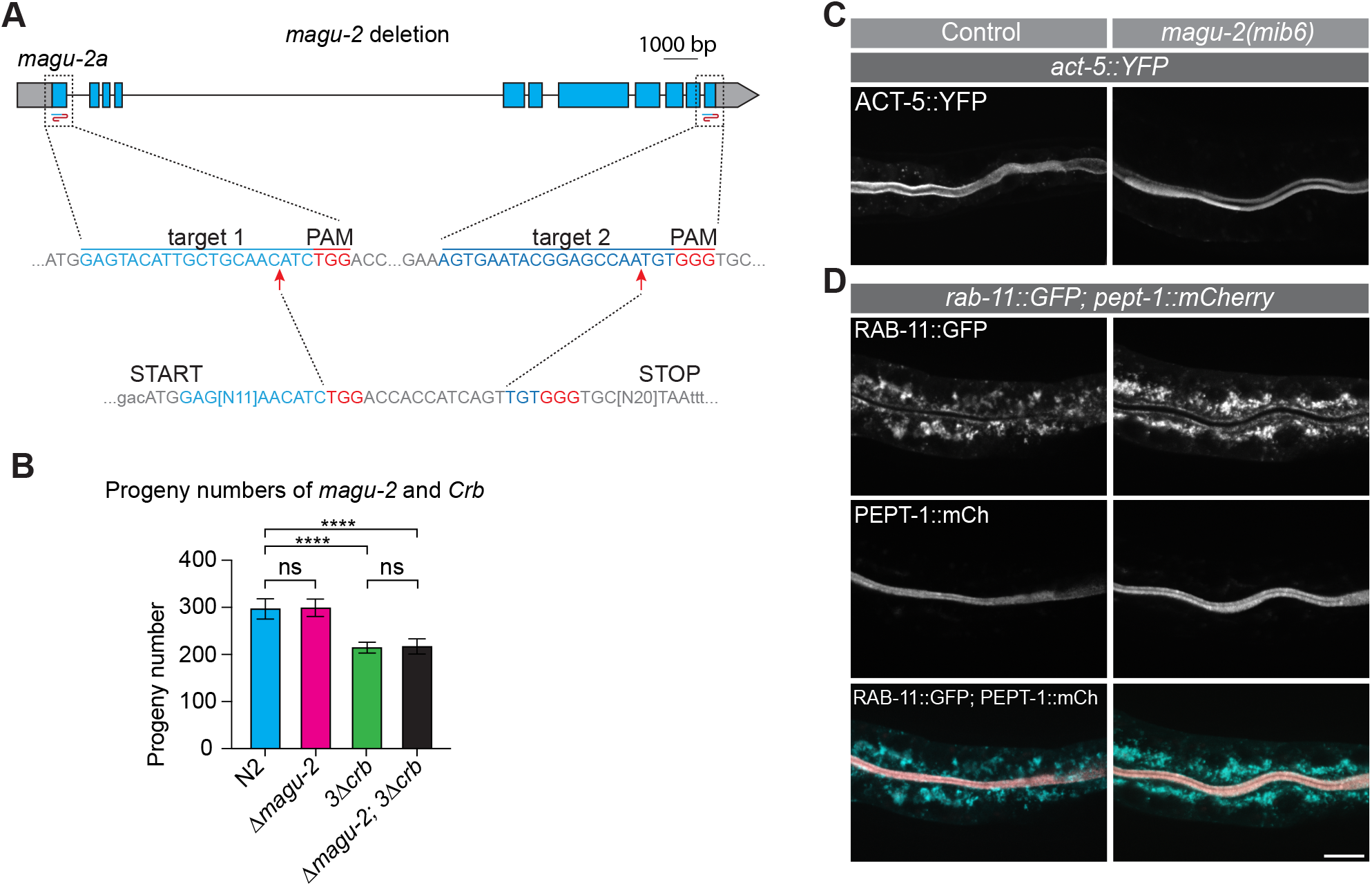
MAGU-2 is not essential for epithelial polarity. **(A)** Top: genomic *magu-2a* gene prediction with sgRNA target sites for the deletion indicated by blue/red ‘S’ shape symbol. Gray boxes indicate untranslated regions. Middle: sgRNA target sites for the deletion indicated in blue text and PAM sites indicated in red text. Bottom: sequence of the obtained deletion. **(B)** Quantifications of brood size. Each symbol represents the progeny of an individual animal (*n* = 7, 7, 7, 10). Error bars are the mean ± s.d. Test of significance: Dunnet’s multiple comparisons test. ns, not significant. **** P ≤ 0.0001. **(C)** Distribution of ACT-5::YFP in control or *magu-2* knock-out (*mib6*) larvae. **(D)** Distribution of RAB-11::GFP and PEPT-1::mCherry in control or *magu-2* knock-out (*mib6*) larvae.

To assess in more detail whether loss of *magu-2* disrupts apical membrane morphology, we examined the localization pattern of the apically enriched intestine-specific actin ACT-5, the apically enriched recycling endosome marker RAB-11, and the apical membrane transporter PEPT-1 in the absence of *magu-2*. ACT-5 and PEPT-1 localized apically in control animals as well as in *magu-2* deletion mutants (Fig, 3C-D). Similarly, RAB-11, which is normally enriched close to the apical lumen in the cytoplasm, maintained the same localization pattern in the absence of MAGU-2 (Fig. 3D). Taken together, these data indicate that, like the *crumbs* genes, *magu-2* is not essential for *C. elegans* development and suggest that it may not be necessary for apical polarity.

### Apical localization of MAGU-2 is dependent on Crumbs

Mammalian Pals1 and *Drosophila* Sdt have been shown to be dependent on Crumbs for their apical localization (Bachmann et al., 2001; Hong et al., 2001; Roh et al., 2002). To determine if the apical localization of MAGU-2 similarly depends on the presence of Crumbs proteins in *C. elegans*, we examined the localization of MAGU-2 in the triple *crumbs* deletion mutant. Absence of the three Crumbs proteins resulted in a complete loss of apical accumulation of MAGU-2 in the embryonic and larval intestine (Fig. 4A), indicating that Crumbs is necessary for apical enrichment of MAGU-2. In some systems studied, Crumbs apical localization is also dependent on Pals1 or Sdt (Almeida et al., 2019; Berger et al., 2007; Hong et al., 2001; Perez-Mockus et al., 2017; Straight et al., 2004; van Rossum et al., 2006). To determine if this reciprocal requirement is conserved in *C. elegans*, we examined the localization of CRB-3 in the *magu-2(mib-6)* deletion. To visualize CRB-3, we used a translational CRB-3::GFP fusion expressed from an integrated extrachromosomal array (Waaijers et al., 2015). In control worms, CRB-3 localizes to the apical side of the intestine in both embryos and larvae. Upon absence of *magu-2* the localization of CRB-3 was unaltered (Fig. 4B), indicating that MAGU-2 does not determine CRB-3 localization in *C. elegans*. Together, these results indicate that the apical recruitment of MAGU-2 depends on Crumbs, while at least the apical localization of CRB-3 is independent of MAGU-2.

**Figure 4.**
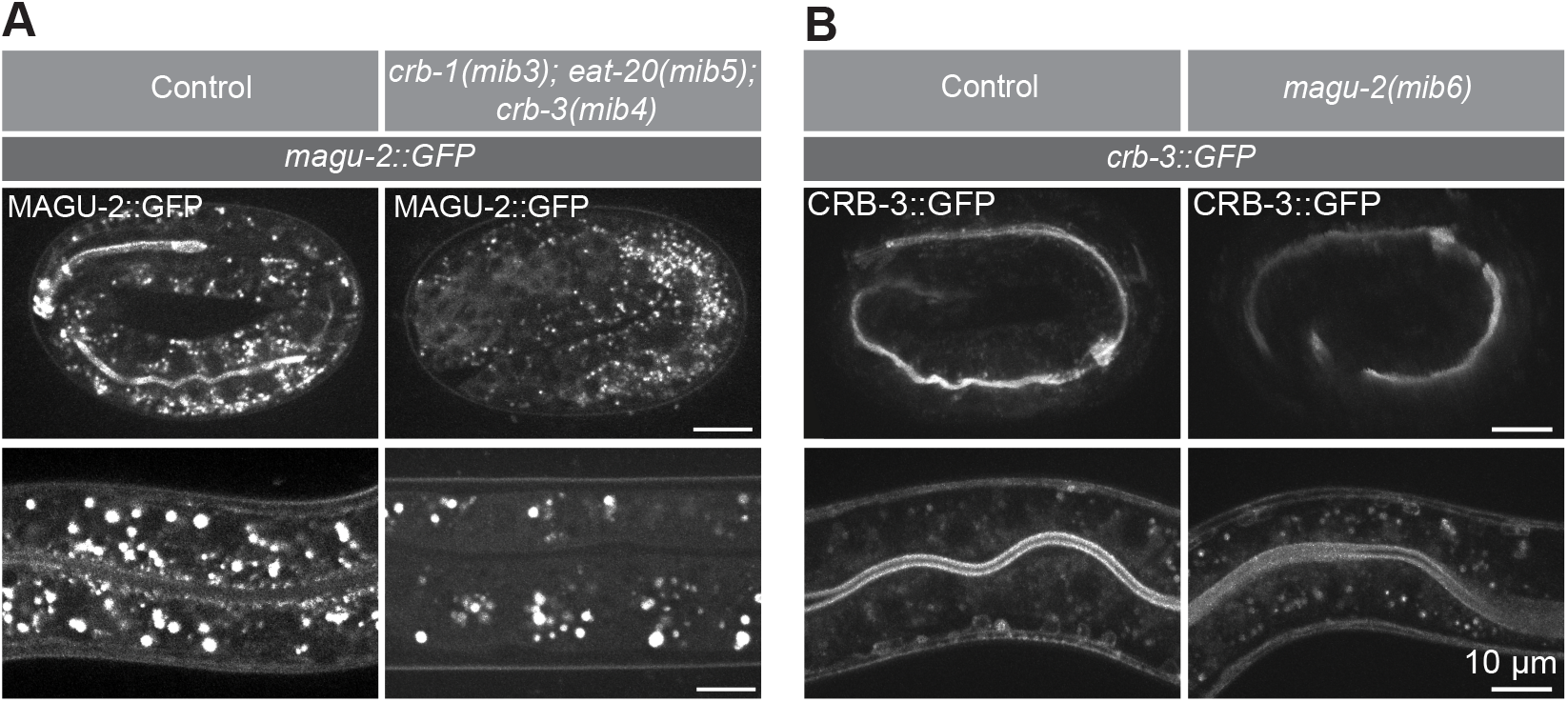
MAGU-2 is dependent on Crumbs for its localization. **(A)** Distribution of MAGU-2::GFP in control or *crb-1(mib3); eat-20(mib5); crb-3(mib4)* two-fold embryos or larvae. **(B)** Distribution of CRB-3::GFP in control or *magu-2* knock-out *(mib6)* three-fold embryos or larvae.

### MAGU-2 interacts with the PATJ ortholog MPZ-1

Having established that the *C. elegans* Crumbs complex contains a Pals1 ortholog, we next investigated the existence of a PATJ ortholog. BLAST searches with the human and *Drosophila* PATJ sequences identified the protein MPZ-1 as the most likely *C. elegans* ortholog. However, reciprocal BLAST using MPZ-1 identified two possible human orthologs, PALS1 and MUPP1, while the most similar *Drosophila* proteins are the basolateral polarity regulators Dlg and Scrib. Therefore, we explored the interaction partners of MAGU-2 in an unbiased fashion, by performing affinity purification of endogenous GFP::MAGU-2 from a mixed-stage population followed by mass-spectrometry. The highest-ranking candidate interacting protein was MPZ-1 (Fig. 5A). These results, together with the homology search, strongly suggest that MPZ-1 is the *C. elegans* ortholog of PATJ. Mammalian PATJ is characterized by the presence of 10 PDZ domains, while *Drosophila* PATJ contains four PDZ domains. Both human and *Drosophila* proteins also contain an N-terminal L27 domain, which interacts with the L27 domain of Pals1 (Roh et al., 2002). Like mammalian PATJ, MPZ-1 contains 10 PDZ domains. However, it lacks a recognizable N-terminal L27 domain. As the L27 domains through which PATJ and Pals1 interact appear to be absent in both MPZ-1 and MAGU-2 (Fig. 1B, 5B), we aimed to identify the MAGU-2 domain responsible for the interaction with MPZ-1. We fused the full length MPZ-1 coding sequence to the DNA binding domain of Gal4 and examined the interaction with the MAGU-2 activation domain fusions described above using Y2H (Fig. 1D). Full length MAGU-2 interacted with MPZ-1, confirming the mass spectrometry result (Fig. 5C). MAGU-2 fragments 2 and 3, which contain the region in which the L27 domain is located in mammalian Pals1 and *Drosophila* Sdt, also interacted with MPZ-1. This suggests that the region between the first PDZ domain and the SH3 domain of MAGU-2 is necessary for the interaction between MAGU-2 and MPZ-1, despite lacking a recognizable L27 domain. Taken together, these data indicate that the interactions between Crumbs, Pals1, and PATJ are conserved in *C. elegans*.

**Figure 5.**
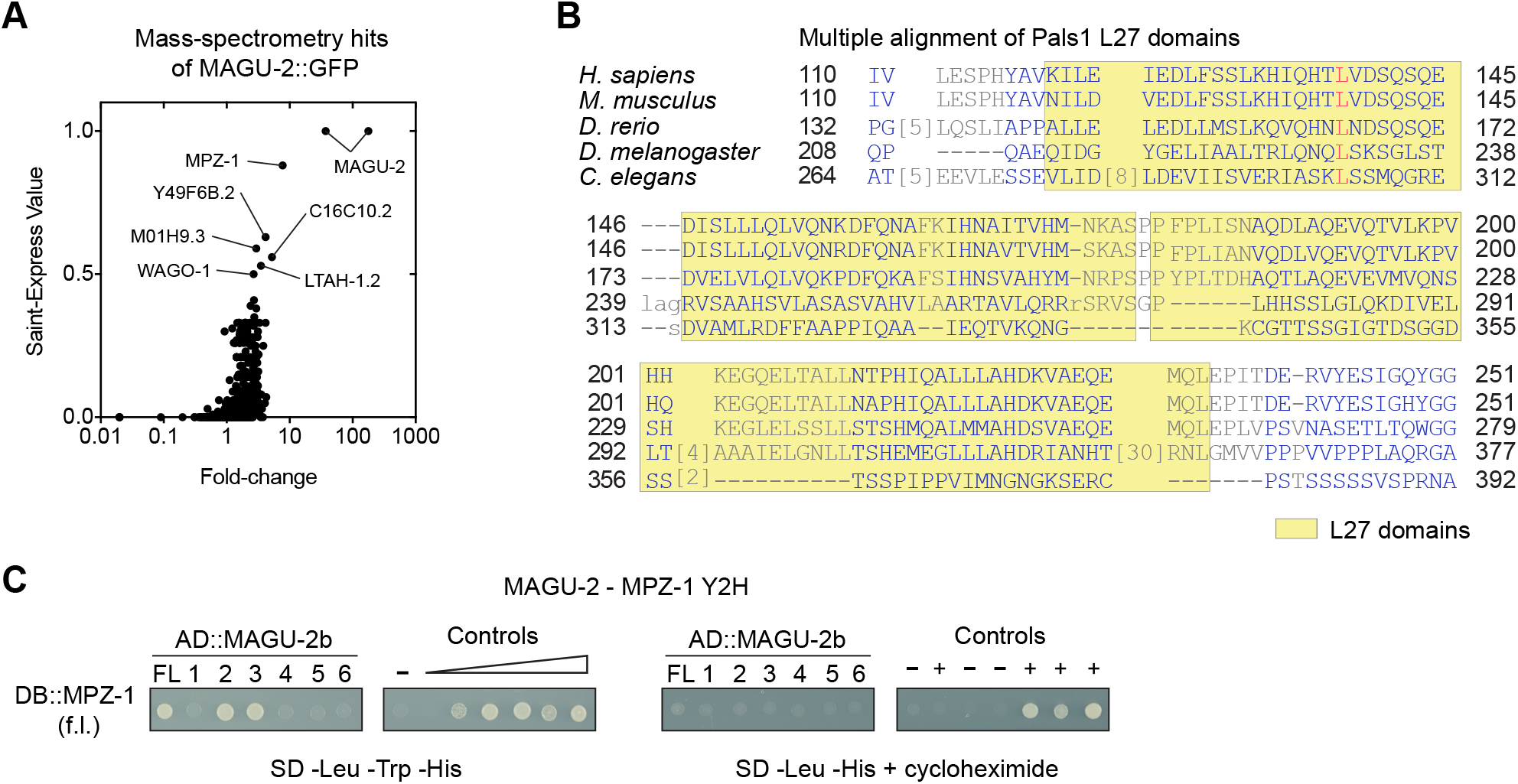
MAGU-2 interacts with MPZ-1. **(A)** Mass spectrometry hits for MAGU-2::GFP pull down plotted as correlation between fold-change score and Saint-Express Value. **(B)** Multiple alignment of the L27 domains in *Homo Sapiens*, *Mus musculus*, *Danio rerio*, *Drosophilamelanogaster* and *Caenorhabditis elegans*. **(C)** Interaction of MAGU-2 and MPZ-1 in the yeast two-hybrid assay. Fragments of MAGU-2.b fused to the Gal4 activation domain were co-expressed with the full length of MPZ-1 fused to the Gal4 DNA binding domain. Growth on the -Leu -Trp -His plate indicates presence of interaction. Lack of growth on -Leu -His + cycloheximide plate shows that DB::MPZ-1 is not self-activating. Controls range from no reporter activation to strong reporter activation.

### Overexpression of Crumbs causes apical membrane domain expansion

Deletion of all three *crumbs* genes demonstrated that the Crumbs complex is not essential in *C. elegans* (Waaijers et al., 2015). However, the effects of Crumbs overexpression, which is known to result in an enlarged apical domain in *Drosophila* (Wodarz et al., 1995), have not been investigated in *C. elegans.* In order to induce overexpression, we expressed EAT-20 and CRB-3 under the control of the intestine specific promoter *elt-2* (McGhee et al., 2009). To determine the effect on apical domain size, we examined the distribution of YFP::ACT-5. Expression levels of EAT-20 and CRB-3 from extrachromosomal arrays varied, but in animals expressing the highest levels of either protein the distribution of ACT-5 indicated an enlarged apical domain which bulged into the cytoplasm of the intestinal cells (Fig. 6). This indicates that overexpression of EAT-20 or CRB-3 results in apical domain expansion, and that the role of the Crumbs proteins in promoting apical domain formation are conserved in *C. elegans*.

**Figure 6.**
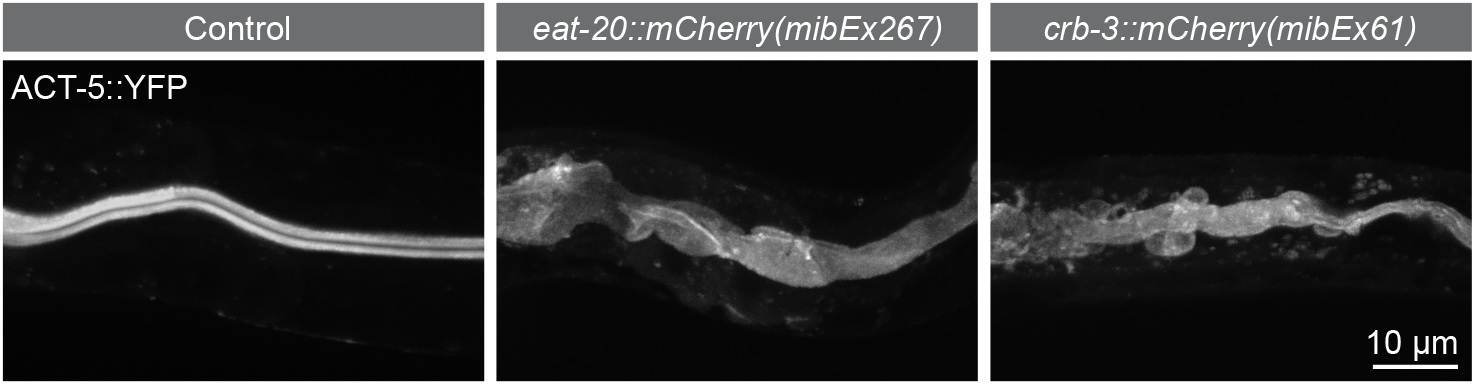
Crumbs overexpression results in apical domain expansion. Distribution of ACT-5::YFP in the intestine of larvae in control animals or upon EAT-20 or CRB-3 overexpression.

## Discussion

Here, we characterized the conservation of composition and function of the Crumbs complex in *C. elegans*. We identified MAGU-2 as the *C. elegans* ortholog of mammalian Pals1 and *Drosophila* Sdt. This conclusion is supported by the sequence conservation, the apical localization of MAGU-2, the dependency of MAGU-2 apical localization on Crumbs proteins, and the interactions with each of the Crumbs orthologs. We also identified MPZ-1 as the ortholog of the Crumbs complex component PATJ, based on sequence conservation and the interaction of MAGU-2 with MPZ-1. Finally, we demonstrate that overexpression of the *C. elegans* Crumbs proteins EAT-20 CRB2 or CRB-3 CRB3 can induce apical membrane expansion in the intestine.

The composition of the Crumbs complex, and the functioning of components besides Crumbs, had not been studied in *C. elegans*. Here, we focused on the homolog of Pals1, *magu-2*, which is a core member of the Crumbs complex in mammalian systems and *Drosophila*. MAGU-2 interacted with *C. elegans* CRB-1, EAT-20, and CRB-3 through its PDZ and SH3 domains, as has been shown for Sdt and Pals1 in *Drosophila* and mammals (Bachmann et al., 2001; Hong et al., 2001; Roh et al., 2002). MAGU-2 was enriched at the apical membrane and relied on CRB-1/EAT-20/CRB-3 for its localization. These results are in accordance with previous observations of Sdt and Pals1, both of which localize apically in a Crumbs dependent manner (Bachmann et al., 2001; Hong et al., 2001; Roh et al., 2002). CRB-3 did not, however, rely on MAGU-2 to become apically enriched. Mammalian Crb3 has been shown to localize apically independently of Pals1 (Straight et al., 2004), whereas Crb1 has been shown to require Pals1 to localize apically (van Rossum et al., 2006). In *Drosophila*, Crumbs has been shown to depend on Sdt in some tissues and developmental stages, such as the embryonic epithelia and mature photoreceptors (Berger et al., 2007; Hong et al., 2001). However in other tissues, such as the posterior midgut and pupal photoreceptors, Crumbs localization is independent of Sdt (Almeida et al., 2019; Perez-Mockus et al., 2017). Thus, the role of Pals1 in localizing Crumbs varies between tissues and organisms. Our efforts here focused on the intestinal localization of CRB-3, hence it remains possible that CRB-1 or EAT-20 depend on MAGU-2 for their apical localization, or that CRB-3 is dependent on MAGU-2 in other tissues.

In addition to the conservation of the interactions with the Crumbs family members, we demonstrated that MAGU-2 interacts with MPZ-1, the likely ortholog of PATJ. Pals1 and PATJ interact through their L27 domains (Feng et al., 2005; Roh et al., 2002), which do not appear to be conserved in *C. elegans* MAGU-2 and MPZ-1. However, the interaction with MPZ-1 was mediated by the region between the first PDZ domain and SH3 domain of MAGU-2, where the L27 domains are localized in Pals1 proteins in other species. This raises the possibility that MAGU-2 contains an L27-like structural fold that is not recognized at the primary amino acid sequence level.

Despite their evolutionary conservation, none of the *C. elegans* Crumbs complex components have essential roles in the development of *C. elegans* (Bossinger et al., 2001; Waaijers et al., 2015). One possible explanation for this difference is that *C. elegans* Crumbs proteins may act redundantly with other proteins. Indeed, CRB-1 was shown to provide a positional cue for junction formation in the intestine upon double depletion of HMP-1 α-catenin and LET-413 Scribble (Segbert et al., 2004), suggesting that redundant mechanisms ensure proper junction formation in the intestine. Further supporting a role for the *C. elegans* Crumbs complex in regulating apical-basal polarization, and similarly to what has been observed in *Drosophila* (Wodarz et al., 1995), we observed that overexpression of either EAT-20 or CRB-3 resulted in apical membrane expansion. Nevertheless, the precise function of the Crumbs proteins remains elusive.

Redundancy in the polarity network has been seen in other systems, such as *Drosophila*, where Crumbs is expressed in all embryonic epithelia derived from the ectoderm but tissues react differently to loss of Crumbs, from disintegration and apoptosis in the epidermis to no apparent defects in the hindgut (Chen et al., 2018; Tepass and Knust, 1990). These findings highlight that in order to understand the different functions of polarity regulators it is important to study them in a range of different systems and organisms. Although it remains uncertain what is the precise role in *C. elegans* of the Crumbs complex, and *magu-2* in particular, our characterization of the Crumbs complex provides further insight into the biology of these evolutionarily conserved components in polarized epithelia.

## Materials and Methods

### Homology searches

Homology searches were performed using BLAST (Altschul et al., 1997) with the input sequences of human Pals1 (Q8N3R9-1) and PatJ (Q9NB04), *Drosophila* Sdt (M9NG38-1) and PatJ-PC (Q8NI35) and *C. elegans* MAGU-2(C01B7.4b.1), MAGU-1 (Q95XW5-1) and MPZ-1 (G5ECZ8-1) were used as input. The following sequences were obtained from UniProt (The UniProt Consortium, 2021) and used for the homology tree: MAGU-1 (Q95XW5), MAGU-2 (Q17549), Pals1(Q8N3R9), and Sdt (Q8WRS3), MPP7 (Q5T2T1) and Metro (A1Z8G0) (Madeira et al., 2019). The homology tree was constructed using Clustal Omega (Sievers et al., 2011) and “Tree of Life” (https://itol.embl.de/itol.cgi) (Letunic and Bork, 2021).

### Yeast two-hybrid analysis

Sequences encoding MAGU-2, CRB-1, EAT-20, CRB-3, and MPZ-1 were PCR amplified from a mixed-stage cDNA library using the primers in Table 1. PCR products were digested with AscI and NotI, and cloned into Gal4-DB vector pMB28 and Gal4-AD vector pMB29, respectively (Koorman et al., 2016). The resulting plasmids were transformed into *Saccharomyces cerevisiae* strains Y8930 (MATα) and Y8800 (MATa) (Yu et al., 2008) using the Te/LiAc transformation method (Schiestl and Gietz, 1989). Diploid yeast was generated by mating, and plated on synthetic defined (SD) medium plates (1) lacking leucine, tryptophan, and histidine and containing 2 mM 3-Amino-1,2,4-triazole (3-AT) to assess the presence of an interaction while reducing background growth; (2) lacking leucine, tryptophan, and adenine to assess the presence of an interaction, and (3) on an SD plate lacking leucine and histidine and containing 1 μg/ml cycloheximide to test for self-activation. Controls of known reporter activation strength and behavior on cycloheximide were also added to all plates.

**Table 1.**
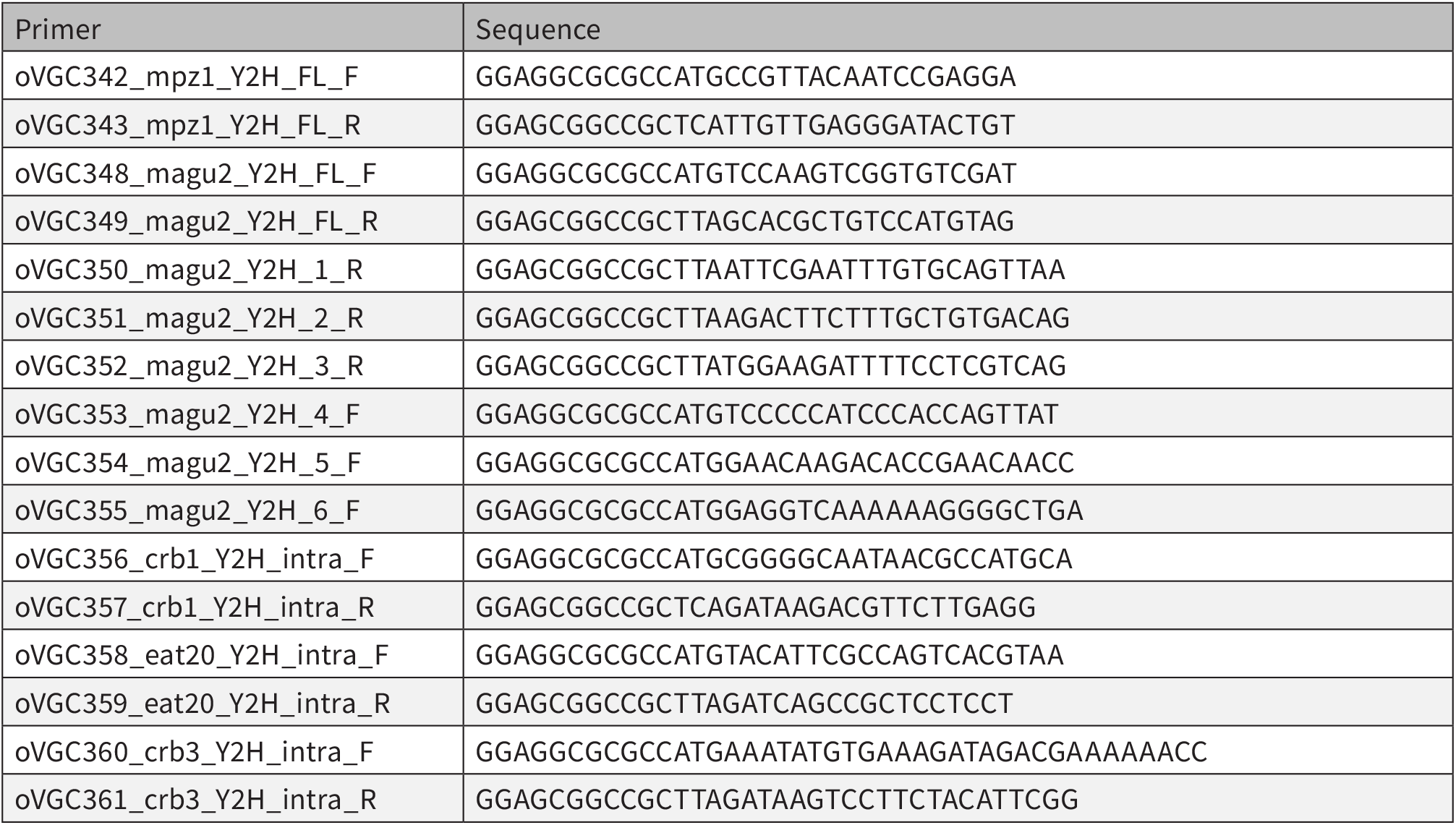
Primers used for Yeast-two-hybrid assay.

### *C. elegans* strains

All *C. elegans* strains used in this study are derived from the N2 Bristol strain, and are listed in Table 2. All strains were maintained at 20 °C on Nematode Growth Medium (NGM) plates seeded with *Escherichiae coli* OP50 bacteria under standard conditions (Brenner, 1974). Table 2 contains a list of all strains used.

**Table 2.** Strain list

| Strain | Genotype | Acknowledgement |
| --- | --- | --- |
| JM125 | *cals108[Pges-1p::YFP::ACT-5]* | James McGhee |
| BOX301 | *magu-2(magu-2::GFP::loxP) V; dlg-1(mib23) X* |  |
| BOX300 | *par-6(mib25[par-6::mCherry-LoxP]) I; magu-2(mib36) V* |  |
| BOX267 | *magu-2(magu-2::GFP::loxP) V* |  |
| BOX42 | *mibls24[crb-3::GFP- Avi, Pmyo-3::mCherry] IV* |  |
| BOX161 | *magu-2(mib6) V* |  |
| BOX164 | *magu-2(mib6) V; crb-1(mib3), eat-20(mib5), crb-3(mib4) X* |  |
| BOX176 | *magu-2(mib6) V; mibls24[crb-3::GFP-2TEV-Avi; Pmyo-3::mCherry] IV* |  |
| BOX198 | *magu-2(mib6) V; cals[Pges-1::YFP::ACT-5]* |  |
| MZE1 | *unc-119(ed3) III; cbgls91[pPept-1:pept-1::DsRed;unc-119(+)]; cbgls98[p-Pept-1:GFP::rab-11.1;unc-119(+)]* | Prof. M. Zerial |
| BOX182 | *magu-2(mib6) V; unc-119(ed3) III; cbgls91[pPept-1:pept-1::DsRed;unc-119(+)]; cbgls98[pPept-1:GFP::rab-11.1;unc-119(+)]* |  |
| BOX283 | *magu-2(magu-2::GFP::loxP) V; crb-1(mib3), eat-20(mib5), crb-3(mib4) X* |  |
| BOX64 | *mibls39[Prps-27::GFP-2xTEV-Avi 10 ng/ul + Prab-3::mCherry 5 ng/ul + lambda DNA 65 ng/ul] I* |  |

### CRISPR/Cas9 genome engineering

The *magu-2::GFP* fusion was done by homology-directed repair of CRISPR/Cas9-induced DNA double-strand breaks in an N2 background. The fusion was generated using plasmid-based expression of Cas9 and sgRNAs. The sequence targeted by the sgRNA was 5’ GTGAATACGGAGCCAATGT 3’. The *magu-2::GFP* repair template was cloned using Sap-Trap assembly into vector pDD379, and the fusion was repaired using a plasmid-based template with 550-600 bp homology arms. A self-excising cassette (SEC) was used for selection (Dickinson et al., 2015). The homology arms included mutations of the sgRNA recognition site to prevent re-cutting after repair. The injection mix was prepared in MilliQ H_2_O and contained 50 ng/μL Peft-3::cas9 (Addgene ID #46168) (Friedland et al., 2013), 65 ng/μL sgRNA-repair-template vector, and 2.5 ng/μL co-injection pharyngeal marker Pmyo-2::tdTomato to aid in visual selection of transgenic strains. Young adult hermaphrodites were injected in the germline using an inverted micro-injection setup (Eppendorf FemtoJet 4x mounted on a Zeiss Axio Observer A.1 equipped with an Eppendorf Transferman 4r). Candidate edited progeny were selected on plates containing 250 ng/μl of hygromycin (Dickinson et al., 2015), and correct genome editing was confirmed by PCR amplification of the edited genomic region. From correctly edited strains, the hygromycin selection cassette was excised by heat shock of L1 larvae at 34 °C for 1 h in a water bath. Correct excision was confirmed by Sanger sequencing (Macrogen Europe) of PCR amplicons encompassing the edited genomic region.

The *magu-2(mib6)* deletion was generated by imprecise repair of a CRISPR/Cas9-induced DNA double-strand breaks and selected using the *dpy-10* co-CRISPR approach (Arribere et al., 2014). The fusion was generated using plasmid-based expression of Cas9 and sgRNAs. Two pairs of sgRNA plasmids were used to target the 5’ and 3’ ends of the *magu-2* open reading frame (ORF), and were generated by ligation of annealed oligo pairs into the *pU6::sgRNA* expression vector pMB70 (Addgene #47942) as previously described (Waaijers et al., 2013). The sequence targeted by the sgRNA were: 5’ GAGTACATTGCTGCAACATC; 5’ GACCACCATCAGTTGCTCCA; 5’ AGTGAATACGGAGCCAATGT; and 5’ AATTCTAATGAAAGTGAATA. The sgRNA targeting the *dpy-10* locus (Arribere et al., 2014) was cloned into the *pU6::sgRNA* expression vector pJJR50 (Addgene #75026) as previously described (Waaijers et al., 2016). The injection mix was prepared in MilliQ H_2_O and contained 60 ng/mL *Peft-3::Cas9* (Addgene #46168), 45 ng/mL each sgRNA plasmid, and 2.5 ng/mL *Pmyo-2::mCherry* as a co-injection marker (pCFJ90, Addgene #19327). Microinjection of adult N2 hermaphrodites was performed as described above. For selection of edited genomes, injected animals were transferred to individual culture plates, incubated for 3-4 days at 20 °C, and 96 non-transgenic F1 animals (wild type, Dpy, or Rol) from 2-3 plates containing high numbers of Dpy and Rol animals were selected and transferred to individual NGM plates. After laying eggs, F1 animals were lysed and genotyped by PCR using two primers flanking the *magu-2* ORF: 5’ to 3’ left primer CATACGCCCAATCATCCGCAC, right primer GATGATGATGGTGTCTCTTCTG. Sanger sequencing was used to determine the precise molecular lesion (Macrogen Europe), and the confirmed knockout strain was back-crossed three times with N2.

### Brood size

Starting at the L4 stage, individual P0 animals were cultured at 20 °C and transferred to a fresh plate every 24 h for 6 days. For each plate, hatched F1 progeny was scored 24 h after removal of the P0.

### Microscopy

Live imaging of *C. elegans* larvae was done by mounting larvae on 5% agarose pads in a 10 mM Tetramisole solution in M9 buffer to induce paralysis. Spinning disk confocal imaging was performed using a Nikon Ti-U microscope driven by MetaMorph Microscopy Automation & Image Analysis Software (Molecular Devices) and equipped with a Yokogawa CSU-X1-M1 confocal head and an Andor iXon DU-885 camera, using 60x or 100x 1.4 NA objectives. All stacks along the z-axis were obtained at 0.25 μm intervals, and all images were analyzed and processed using ImageJ (FIJI) and Adobe Photoshop.

### GFP pull-down of MAGU-2::GFP

Animals endogenously expressing GFP-tagged MAGU-2 or control animals expressing an integrated GFP transgene (Waaijers et al., 2016) were grown on 6–8 9 cm NGM plates until starvation, to enrich for L1 animals. Animals were then transferred into 250 ml of S-Medium supplemented with 1% Penn/Strep (Life Technologies), 0.1% nystatin (Sigma) and OP50 bacteria obtained from the growth of a 400 ml culture. Animals were grown at 20 °C at low shaking for 96 hours and were harvested and cleaned using a sucrose gradient, as previously described (Waaijers et al., 2016) with one exception being the inclusion of MgSO4 in the M9 medium. Worms were distributed into 15 ml TPX tubes (Diagenode) to reach 200–400 μl worm pellet per tube, and were washed with lysis buffer (25mM Tris-HCl pH 7.5, 150mM NaCl, 1mM EDTA, 0.5% IGEPAL CA-630, 1X Complete Protease Inhibitor Cocktail (Roche)). The liquid was removed, and the sample was flash frozen in liquid nitrogen for storage at −80 °C. To lyse the worms, tubes were thawed on ice and ice-cold lysis buffer was added to reach a total volume of 2 ml. Tubes were sonicated for 10 mins (sonication cycle: 30 sec ON, 30 sec OFF) at 4 °C in a Bioruptor ultrasonication bath (Diagenode) at high energy setting. After lysis, lysates were cleared by centrifugation and protein levels were measured using the Bradford BCA assay (Thermo Scientific). Immunoprecipitation was performed using GFP-Trap Magnetic Agarose beads (Chromotek) according to manufacturer’s protocol, using 25 μl of beads per sample. To prep the beads, they were first equilibrated in wash buffer (10 mM Tris/Cl pH 7.5, 150 mM NaCl, 0.5 mM EDTA, 0.1% IGEPAL CA-630), blocked with 1% BSA for 1 hour, then washed 4 times with wash buffer. Next, lysate was added to the beads and they were incubated for 1 hour tumbling end-over-end. Lysate was then removed, and the beads were washed 4 times in wash buffer. After the final wash step, all liquid was removed, and the beads were flash frozen with liquid nitrogen. The experiment was performed in triplicate (biological replicates) and processed on independent days.

### Mass spectrometry analysis for MAGU-2

After affinity purification anti-GFP beads were resuspended in 15 μl of 4× Laemmli sample buffer (Biorad), boiled at 99 °C for 10 min and supernatants were loaded on 4–12% Criterion XT Bis–Tris precast gel (Biorad). The gel was fixed with 40% methanol and 10% acetic acid and then stained for 1 hour using colloidal coomassie dye G-250 (Gel Code Blue Stain, Thermo Scientific). Each lane from the gel was cut and placed in 1.5 ml tubes. Samples were then washed with 250 μl of water, followed by 15 min dehydration in acetonitrile. Proteins were reduced (10mM DTT, 1 hour at 56 °C), dehydrated and alkylated (55mM iodoacetamide, 1 hour in the dark). After two rounds of dehydration, trypsin was added to the samples and incubated overnight at 37 °C. Peptides were extracted with acetonitrile, dried down and reconstituted in 10% formic acid prior to MS analysis.

Samples were analyzed on an Orbitrap Q-Exactive mass spectrometer (Thermo Fisher Scientific) coupled to an Agilent 1290 Infinity LC (Agilent Technologies). Peptides were loaded onto a trap column (Reprosil pur C18, Dr. Maisch, 100 μm × 2 cm, 3 μm; constructed in-house) with solvent A (0.1% formic acid in water) at a maximum pressure of 800 bar and chromatographically separated over the analytical column (Poroshell 120 EC C18, Agilent Technologies, 100 μm x 50 cm, 2.7 μm) using 90 min linear gradient from 7% to 30% solvent B (0.1% formic acid in acetonitrile) at a flow rate of 150 nl min^−1^. The mass spectrometers were used in a data-dependent mode, which automatically switched between MS and MS/MS. After a survey scan from 375 to 1600m/z the 10 most abundant peptides were subjected to HCD fragmentation. MS spectra were acquired with a resolution > 30,000, whereas MS2 with a resolution > 17,500. Raw data files were converted to mgf files using Proteome Discoverer 1.4 software (Thermo Fisher Scientific). Database search was performed using the *C. elegans* database and Mascot (version 2.5.1, Matrix Science, UK) as the search engine. Carbamidomethylation of cysteines was set as a fixed modification and oxidation of methionine was set as a variable modification. Trypsin was set as cleavage specificity, allowing a maximum of two missed cleavages. Data filtering was performed using a percolator, resulting in 1% false discovery rate (FDR). Additional filters were search engine rank 1 and mascot ion score > 20.

Crapome (Mellacheruvu et al., 2013) was used to analyze MAGU-2 interacting proteins in three biological replicas, using proteins identified in the GFP pull downs, as well as in pull downs of eGFP::PAR-3 (BOX290), GFP::P-KC-3 (KK1228) and DLG-1::eGFP (BOX260) as control. Significance analysis of interactome (SAINT) score (Choi et al., 2011) and simpler fold-change (FC) calculations FC-A and FC-B were derived from the Crapome analysis by averaging the spectral counts across the controls. FC-A averages the counts across all controls while the more stringent FC-B takes the average of the top 3 highest spectral counts for the abundance estimate.

## Acknowledgements

We thank members of the S van den Heuvel and M Boxem groups for helpful discussions. We also thank Wormbase (Harris et al., 2020) and the Biology Imaging Center, Faculty of Sciences, Department of Biology, Utrecht University. Some strains were provided by the Caenorhabditis Genetics Center, which is funded by NIH Office of Research Infrastructure Programs (P40 OD010440). This work was supported by the Netherlands Organization for Scientific Research (NWO)-ALW Open Program 824.14.021 and NWO-VICI 016.VICI.170.165 grants to M Boxem, and the European Union’s Horizon 2020 research and innovation programme under the Marie Skłodowska-Curie grant agreement No. 675407 – PolarNet. This research was also part of the Netherlands X-omics Initiative and partially funded by NWO (Project 184.034.019).

## Author contributions

Conceptualization, V.G.C., J.J.R., and M.B.; Formal Analysis, V.G.C., J.J.R., J.K., and H.v.B.; Investigation, V.G.C., J.J.R., J.K., H.v.B., and R.S.; Resources, M.B.; Data Curation, V.G.C.; Writing – Original Draft, V.G.C.; Writing – Review & Editing, V.G.C. and M.B.; Visualization, V.G.C. and M.B.; Supervision, V.G.C. and M.B.; Project Administration, M.B.; Funding Acquisition, M.B.

